# Sexual dimorphism in molecular profiles of resting human skeletal muscle and the response to acute exercise and endurance training

**DOI:** 10.1101/2024.12.12.627717

**Authors:** Simon Dreher, Thomas Goj, Christine von Toerne, Miriam Hoene, Martin Irmler, Meriem Ouni, Markus Jähnert, Johannes Beckers, Martin Hrabe de Angelis, Andreas Peter, Anja Moller, Andreas L. Birkenfeld, Annette Schürmann, Stefanie M. Hauck, Cora Weigert

## Abstract

Biological sex has a strong impact on skeletal muscle metabolism and performance. By a comprehensive investigation of epigenetic, transcriptomic and proteomic differences between female and male skeletal muscle of untrained subjects we provide a molecular basis for the sexual dimorphism of glucose and lipid metabolism. The sex-specific multi-OMICs profiles indicate higher degree of glucose turnover and higher abundance of fast-twitch fibers in males and high degree of lipid handling in females. Eight-week endurance training equalized initial differences toward an endurance-trained proteomic profile in both sexes. The untrained muscle of females was more resistant to an acute exercise challenge since stress-responsive transcripts were predominantly upregulated in males.

In myotubes from the same donors, transcriptomic differences were hardly conserved, but could be partially restored by treatment with sex hormones. In conclusion, after only 8 weeks training mitigates deeply rooted sex-specific molecular profiles in skeletal muscle toward a common metabolically beneficial response.

## Introduction

The skeletal muscle comprises 35 – 45% of total body weight in healthy non-obese females and males and is of significant importance for the regulation of glucose and fatty acid metabolism. It is responsible for 85% of insulin-dependent glucose uptake, and glucose uptake can increase up to 50-fold during intense muscular work ^1^. The impact of biological sex on the skeletal muscle metabolism in health and disease, including physical activity and prevention of type 2 diabetes, has been neglected for decades. Historically, for multiple reasons, biomedical research has been centered on male physiology. Pre-clinical studies and clinical trials that include women did not analyze the results by sex. This perception is undergoing significant changes, since it has been recognized that the biological sex has a clear impact on the regulation of peripheral metabolism and insulin sensitivity ^2^, with implications for the pathophysiology and clinical manifestation of metabolic diseases and response to treatment ^3–5^. Hyperinsulinemic-euglycemic clamp data which consider differences in body weight or lean mass indicate higher glucose disposal in women, driven by higher insulin-induced glucose uptake in skeletal muscle ^6–9^. Several studies also reported greater clearance of triglycerides from VLDL in women ^10–13^, driven again by greater triglyceride extraction by skeletal muscle ^14^. Together with the higher plasma fatty acid availability for skeletal muscle repeatedly reported in women ^15^, this may cause the higher intramyocellular lipid content in female skeletal muscle ^16,17^. Notably, not only intramyocellular triglyceride (IMTG) storage but also IMTG lipolysis is suggested to be higher in women than in men ^18^.

Beyond that, sex as a biological variable influences exercise metabolism and potentially the adaptation to regular training. It has been reported that during exercise, females utilize more fat, and males more carbohydrates ^19–22^. In contrast, after exercise, females showed elevated glucose oxidation and males elevated lipid oxidation ^15,23^. The response to training may also differ since compared to females, males were reported to have greater increase in VO_2_peak, muscle mass and strength in response to exercise despite already higher levels at baseline ^24–26^.

A different content of type 1 and 2 muscle fibers in female and male skeletal muscle may explain at least some of the differences in metabolism ^27^. Staining for fiber type-specific myosin heavy chains or enzymes showed that type I fibers cover a greater muscle area in women than in men ^18,27–30^. Human skeletal muscles contain three major fiber types, slow type 1, and fast type 2A and 2X, which are best characterized by the dominant abundance of MYH7, MYH2 and MYH1 proteins, respectively, and a fiber type-specific pattern of structural proteins and metabolic enzymes ^31^. The oxidative type 1 fibers have a higher abundance of mitochondrial proteins and proteins relevant for fatty acid handling, whereas the fast fibers have a higher content of glycolytic enzymes. Thus, higher percentage of type 1 fibers in female muscle would support a higher capacity of lipid storage and turnover as well as substrate oxidation. In conclusion, sex-specific differences in glucose and fatty acid handling in skeletal muscle of women and men exist, with potential implications for prevention and treatment of metabolic diseases.

The molecular basis for the sexual dimorphism in skeletal muscle metabolism is incompletely understood and the impact of sex on the response of the transcriptome and proteome to exercise need further elucidation. Investigations in the field mainly covered transcriptional and epigenetic differences ^32–34^, or compared the skeletal muscle proteome of both sexes in an untrained and a trained group ^35^. In our study, we aim to provide a yet missing comprehensive picture of molecular differences between female and male skeletal muscle to explain the observed differences in glucose and lipid metabolism. Using skeletal muscle biopsies obtained during an 8-week endurance training intervention ^36,37^, we applied a multi-omics approach employing DNA methylation, transcriptomics and proteomics. We studied not only the differences at baseline, but also sex-based differences in the response to acute exercise and after the 8-week endurance training. Moreover, we investigated whether sex-specific differences were conserved in the myotubes generated from the myogenic satellite cells of the same donors and whether treatment with sex hormones can mimic a sex-specific transcriptional regulation identified in vivo.

## Methods

### Study participants

Study design and participants were described previously ^36^. Inclusion criteria for the study were sedentary lifestyle (<120 minutes of physical activity per week), BMI >27 kg/m^2^, but absence of manifest diabetes. All but one female were premenopausal. Two females used oral contraceptives. Baseline muscle biopsies and training was not synchronized with the menstrual cycle. All participants gave written informed consent and the study protocol was approved by the ethics committee of the University of Tübingen and was in accordance with the declaration of Helsinki. The study was registered at Clinicaltrials.gov as trial number NCT03151590.

### Study design

Participants performed one hour of supervised endurance training three times per week for 8 weeks, consisting of 30 minutes of cycling on an ergometer and 30 minutes of walking on a treadmill. Before and after the training period, all participants underwent maximal spiroergometry as an incremental cycling test using an electromagnetically braked bicycle ergometer, to determine the individual VO_2_peak. The test was terminated at volitional exhaustion or muscular fatigue. Peak VO_2_ was defined as the mean VO_2_ over the last 20 s before the cessation of exercise and was assessed by metabolic gas analysis. The training intensity was individually set at 80% of the VO_2_peak determined before the intervention and was not changed throughout the training period. Training intensity was controlled by heart rate based on predetermined 80% of the VO_2_peak and individually set. For anthropometric data of the included 16 females and 9 males, see Tab. 1. Body fat mass and distribution were measured by magnetic resonance imaging ^38^. Metabolic and fitness parameters were assessed as described in ^36^. The insulin sensitivity index was estimated from the oGTT by the method of Matsuda and DeFronzo (ISI) ^39^.

**Table 1.** Anthropometric data of study participants.

|  | Baseline |  |  | Trained |  |  | Fold change |  |  |
| --- | --- | --- | --- | --- | --- | --- | --- | --- | --- |
|  | Female | Male | p-Value | Female | Male | p-Value | Female | Male | p-Value |
| Sex | 16 | 9 |  |  |  |  |  |  |  |
| Height [cm] | 164 | 181 | **<0.001** |  |  |  |  |  |  |
|  | ± 4.13 | ± 6.23 |  |  |  |  |  |  |  |
| Weight [kg] | 86.4 | 101 | **0.020** |  |  |  |  |  |  |
|  | ± 14.6 | ± 13.8 |  |  |  |  |  |  |  |
| Age [Years] | 27.9 | 33.2 | **0.005** |  |  |  |  |  |  |
|  | ± 8.79 | ± 5.59 |  |  |  |  |  |  |  |
| BMI | 31.9 | 30.6 | 0.479 | 31.7 | 30.2 | 0.350 | 0.99 | 0.99 | 0.567 |
|  | ± 4.37 | ± 3.63 |  | ± 4.45 | ± 3.92 |  | ± 0.02 | ± 0.02 |  |
| Waist to hip | 0.86 | 0.94 | **0.001** | 0.86 | 0.93 | **0.001** | 1 | 0.99 | 0.404 |
|  | ± 0.04 | ± 0.04 |  | ± 0.05 | ± 0.04 |  | ± 0.02 | ± 0.02 |  |
| TAT [L] | 42.4 | 35.4 | **0.044** | 41.7 | 34.2 | **0.039** | 0.98 | 0.97 | 0.508 |
|  | ± 11.0 | ± 9.17 |  | ± 11.1 | ± 8.95 |  | ± 0.03 | ± 0.05 |  |
| SCAT [L] | 16.4 | 13.0 | **0.044** | 15.8 | 12.4 | 0.075 | 0.96 | 0.95 | 0.819 |
|  | ± 5.47 | ± 4.58 |  | ± 5.85 | ± 4.30 |  | ± 0.06 | ± 0.07 |  |
| VAT [L] | 2.69 | 4.72 | **0.004** | 2.62 | 4.42 | **0.006** | 0.98 | 0.94 | 0.296 |
|  | ± 1.14 | ± 1.42 |  | ± 1.14 | ± 1.36 |  | ± 0.16 | ± 0.05 |  |
| AT-L.Ex [L] | 17.9 | 13.1 | **0.008** | 17.9 | 12.5 | **0.003** | 1 | 0.95 | **0.010** |
|  | ± 4.00 | ± 3.53 |  | ± 3.86 | ± 3.42 |  | ± 0.05 | ± 0.03 |  |
| AT femur [L] | 516 | 361 | **0.001** | 495 | 350 | **0.003** | 0.96 | 0.98 | 0.932 |
|  | ± 104 | ± 77.7 |  | ± 109 | ± 68.7 |  | ± 0.08 | ± 0.07 |  |
| NAT-L.Ex [L] | 15.5 | 22.0 | **<0.001** | 16.0 | 22.1 | **<0.001** | 1.03 | 1.01 | **0.048** |
|  | ± 2.16 | ± 3.00 |  | ± 1.93 | ± 2.99 |  | ± 0.04 | ± 0.02 |  |
| IAT/BW | 1.06 | 1.18 | 0.135 | 1.28 | 1.42 | 0.152 | 1.21 | 1.21 | 0.994 |
|  | ± 0.23 | ± 0.13 |  | ± 0.27 | ± 0.19 |  | ± 0.12 | ± 0.12 |  |
| Systolic BP [mm Hg] | 130 | 139 | 0.184 | 124 | 141 | **0.007** | 0.96 | 1.02 | 0.107 |
|  | ± 11.6 | ± 14.8 |  | ± 12.9 | ± 12.2 |  | ± 0.13 | ± 0.13 |  |
| Diastolic BP [mm Hg] | 87.6 | 89.8 | 0.722 | 80.2 | 89.2 | **0.038** | 0.92 | 1.02 | 0.154 |
|  | ± 9.00 | ± 15.3 |  | ± 11.4 | ± 7.88 |  | ± 0.12 | ± 0.16 |  |
| Glucose fast. [mmol/L] | 5.01 | 5.28 | 0.121 | 4.87 | 5.28 | **0.016** | 0.97 | 1 | 0.274 |
|  | ± 0.33 | ± 0.40 |  | ± 0.28 | ± 0.37 |  | ± 0.05 | ± 0.06 |  |
| Glucose 120 [mmol/L] | 5.65 | 5.73 | 0.879 | 5.33 | 5.84 | 0.932 | 0.97 | 1.06 | 0.821 |
|  | ± 1.23 | ± 1.13 |  | ± 0.89 | ± 2.23 |  | ± 0.2 | ± 0.49 |  |
| Insulin fast. [pmol/L] | 110 | 95.6 | 0.396 | 106 | 89.7 | 0.307 | 1.06 | 0.98 | 0.561 |
|  | ± 44.5 | ± 34.4 |  | ± 35.8 | ± 35.5 |  | ± 0.38 | ± 0.28 |  |
| Insulin 120 [pmol/L] | 626 | 401 | 0.533 | 496 | 387 | 0.322 | 0.85 | 1.11 | 0.671 |
|  | ± 463 | ± 186 |  | ± 322 | ± 359 |  | ± 0.32 | ± 1.2 |  |
| ISI<sub>MATS</sub> | 8.22 | 9.58 | 0.799 | 9.00 | 9.63 | 0.755 | 1.14 | 1.1 | 0.843 |
|  | ± 3.38 | ± 6.80 |  | ± 4.10 | ± 5.02 |  | ± 0.37 | ± 0.24 |  |
| HbA1C [mmol/mol] | 33.9 | 35.0 | 0.286 | 33.4 | 33.8 | 0.753 | 0.99 | 0.97 | 0.451 |
|  | ± 2.64 | ± 2.16 |  | ± 2.42 | ± 2.70 |  | ± 0.05 | ± 0.05 |  |
| HbA1C [%] | 5.25 | 5.35 | 0.286 | 5.21 | 5.24 | 0.753 | 0.99 | 0.98 | 0.451 |
|  | ± 0.24 | ± 0.20 |  | ± 0.22 | ± 0.25 |  | ± 0.03 | ± 0.03 |  |
| VO<sub>2</sub>peak/BW | 24.1 | 26.6 | **0.048** | 26.0 | 28.9 | 0.130 | 1.09 | 1.09 | 0.915 |
|  | ± 4.26 | ± 2.29 |  | ± 4.58 | ± 3.78 |  | ± 0.13 | ± 0.15 |  |
| Performance [W] | 101 | 144 | **<0.001** | 113 | 150 | **<0.001** | 1.15 | 1.05 | 0.291 |
|  | ± 20.2 | ± 19.9 |  | ± 13.7 | ± 18.5 |  | ± 0.19 | ± 0.07 |  |

BMI: body mass index; TAT: total adipose tissue; SCAT: subcutaneous adipose tissue; VAT: visceral adipose tissue; AT: adipose tissue; L.Ex: lower extremities; NAT: non-adipose tissue; IAT: individual anaerobic threshold; BW: body weight; BP: blood pressure: fast.: fasted; ISI<sub>MATS</sub>: insulin sensitivity index estimated by the method of Matsuda and DeFronzo<sup>39</sup>; HbA1C: hemoglobin A1C; VO<sub>2</sub>peak: peak oxygen consumption

### Biopsy collection

Skeletal muscle biopsies were obtained from the lateral part of vastus lateralis muscle. After local anesthesia, skin, fat tissue, fascia, and the muscle epimysium were cut under sterile conditions using a scalpel, and a piece of muscle was removed using a Bergström needle (Pelomi Medical, Albertslund, Denmark) with suction. Biopsies were taken 8 days before (baseline, B) and 5 days after (trained, T) the 8-week intervention in a resting state 60 min after the end of an OGTT as well as 60 minutes after the first ergometer exercise bout (acute, A) (Figure S 1). 45 min before the acute exercise bout, participants received a defined breakfast to account for the glucose-induced hormonal changes after the OGTT in the rested state biopsies. All biopsies were collected at 11:00 am ± 30 minutes.

### Proteomics sample processing

Frozen biopsies were lysed in 200µl pre-cooled sodium deoxycholate (SDC) lysis buffer containing 4% (w/v) SDC and 100 mM Tris-HCl (pH 8.5) in tubes filled with 1.4 mm ceramic beads (0.5 mL CK14 soft tissue homogenizing, Bertin, Montigny-le-Bretonneux, France). The samples were homogenized by shaking twice for 20 seconds at 5500 cycles per minute separated by a 5 second break in a Precellys 24 homogenizer (Bertin, Montigny-le-Bretonneux, France). After cell disruption, the mixture was boiled for 5 min at 95 °C. Following snap-freezing on dry ice, samples were sonicated for 5 intervals with 1 sec pulses, 5 seconds off and an intensity of 80 (Probe Sonicator EppiShear, ActiveMotif, Carlsbad, USA). A BCA assay (Thermo Fisher Scientific, Waltham, MA, USA) was performed and 800 µg total protein per sample were used as starting material. Reduction and alkylation were performed in a one pot reaction at 45 °C for 10 minutes using 20 mM chloracetamide (Sigma Alrich, Steinheim, Germany) and 5 mM tris(2-carboxymethyl)phosphine (Bond-BreakerTM TCEP, Thermo Fisher Scientific). In-solution digestion was performed overnight in a ThermoMixer at 2000 rpm at 37 °C with a protein-to enzyme ratio of 1 to 100 for both LysC (Wako, Osaka, Japan) and Trypsin (Promega, Madison, WI, USA). Tryptic peptides were acidified with 1% trifluoroacetic acid (TFA). 40 µg of the peptide solution was cleared by centrifugation and loaded onto activated (washed first with 100% acetonitrile followed by 30% methanol in 1% TFA and finally 0.2% TFA) three-layer styrene divinylbenzene-reversed phase sulfonated STAGE tips (SDB-RPS 3 M Empore, CDS Analytical, Oxford, PA, USA) as described ^40^ with minor adjustments: The STAGE tips containing peptides were first washed with 100 µl 1% TFA in ethyl acetate, 100 µl 1% TFA in isopropanol and 150 µl 0.2% TFA. The peptides were eluted with 5% NH4OH in 80% acetonitrile. Samples were dried completely in a SpeedVac (Concentrator plus, Eppendorf, Hamburg, Germany) for 40 minutes at 45 °C and stored at - 20 °C until MS measurement.

### Mass spectrometric (MS) measurements

LC-MSMS analysis was performed in data-dependent acquisition (DDA) mode. MS data were acquired on a Q-Exactive HF-X mass spectrometer (Thermo Fisher Scientific) online coupled to a nano-RSLC (Ultimate 3000 RSLC; Dionex). Tryptic peptides were automatically loaded onto a C18 trap column (300 µm inner diameter (ID) × 5 mm, Acclaim PepMap100 C18, 5 µm, 100 Å, Thermo Fischer Scientific) at flow rate of 30 µl/min. Peptides were further separated on a C18 reversed phase analytical column (nanoEase MZ HSS T3 Column, 100Å, 1.8 µm, 75 µm x 250 mm, Waters, Milford, MA, USA) at 250 nl/min flow rate in a 95-minutes non-linear acetonitrile gradient from 3% to 40% in 0.1% formic acid. The high-resolution (60000 full width at half-maximum) MS spectrum was acquired with a mass range from 300 to 1500 m/z with automatic gain control target set to 3 x 10^6^ and a maximum of 30 ms injection time. From the MS prescan, the 15 most abundant peptide ions were selected for fragmentation (MSMS) if at least doubly charged, with a dynamic exclusion of 30 seconds. MSMS spectra were recorded at 15000 resolution with automatic gain control target set to 1 x 105 and a maximum of 50 ms injection time. The normalized collision energy was 28, and the spectra were recorded in profile mode.

### Proteome data processing – Protein identification and label-free quantification

Proteome Discoverer (PD) software (Thermo Fisher Scientific, version 2.4.1.15) was used for peptide and protein identification via a database search (Sequest HT search engine) against the Swissprot human data base (Release 2020_02, 20349 sequences in PD), considering full tryptic specificity, allowing for up to two missed tryptic cleavage sites, precursor mass tolerance 10 ppm, fragment mass tolerance 0.02 Da. Carbamidomethylation of Cys was set as a static modification. Dynamic modifications included deamidation of Asn or Gln, oxidation of Met, and a combination of Met loss with acetylation on the protein N-terminus. Percolator was used to validate peptide spectrum matches and peptides, accepting only the top-scoring hit for each spectrum (high confidence), FDR cutoff values of FDR <1%, and a posterior error probability of <0.01. The final list of proteins complied with the strict parsimony principle.

Protein abundances for quantification were based on peak intensity values of proteotypic peptides. Normalization was performed on total peptide amount to account for sample loading errors. The protein abundances were calculated summing up the single abundance values for admissible peptides. Only proteins quantified in at least 90% of the samples from at least one experimental group were included in the statistical analysis. Missing values (2607, 1.43 %) were imputed using a downshift of 1.5 from the mean and a width of 0.5 from the SD.

### Transcriptomic analysis

Total RNA isolation from snap-frozen biopsies and microarray analysis was described in ^41^. Tissues were homogenized using a TissueLyser II (Qiagen) and RNA was isolated with the miRNeasy Kit (Qiagen) including DNAse digestion. Only high-quality RNA (RNA integrity number > 7, Agilent 2100 Bioanalyzer) was used for microarray analysis. Total RNA was amplified using the WT PLUS Reagent Kit (Thermo Fisher Scientific Inc., Waltham, USA). Amplified cDNA was hybridized on Human Clariom S arrays (Thermo Fisher Scientific). Staining and scanning (GeneChip Scanner 3000 7G) was done according to manufacturer’s instructions. Transcriptome Analysis Console (TAC; version 4.0.0.25; Thermo Fisher)) was used for quality control and to obtain annotated normalized SST-RMA gene-level data. Statistical analyses were performed with R3.6.3/Rstudio ^42^.

### DNA methylation analysis

Genomic DNA was isolated from snap-frozen biopsies at baseline using the Invisorb Genomic DNA Kit II (STRATEC Molecular GmbH, Berlin, Germany). The bisulfite treatment and hybridization of the DNA samples were carried out by Life & Brain (Bonn, Germany). DNA methylation levels were determined by the Infinium® MethylationEPIC BeadChip version 1. The data were processed using R (v.4) packages “meffil” (v.1.3.1) and “ChAMP” (v. 2.24.0) as described earlier ^43,44^.

### Cell Culture

Primary human myoblasts were obtained from the muscle biopsies as described previously^45^. Only baseline biopsies were used. Myoblast isolation and enrichment of CD56+ myoblasts by magnetic bead cell sorting are described in ^46^. In brief, satellite cells were released by collagenase digestion and seeded on 15-cm dishes coated with GelTrex™ (Thermo Fisher Scientific, Germany). After two rounds of proliferation in cloning medium (α-MEM:Ham’s F- 12 (1:1), 20% (v/v) FBS, 1% (v/v) chicken extract, 2 mM L-glutamine, 100 units/ml penicillin, 100 µg/ml streptomycin, 0.5 µg/ml amphotericin B), CD56-positive myoblasts were enriched (>90%) using MACS microbeads and LS columns (Milteny Biotech, Germany), according to the manufacturer’s protocol, with a 30-min incubation. They were then stored in the gaseous phase of liquid nitrogen. Cell culture surfaces were prepared with a non-gelling thin-layer GelTrex™ coating. Myoblasts (passage 3 after isolation, passage 1 after enrichment) were proliferated in cloning media until 90% confluency. Myotube differentiation was induced on day 0 as described previously ^47^ and maintained for 7 days in fusion media (α- MEM, 2 mM L-glutamine, 50 µM palmitate, 50 µM oleate (complexed to BSA with a final BSA concentration of 1.6 mg/ml in medium), 100 µM carnitine) with supplementation of 25 ng/ml (13.16 nmol/l) IGF1 (human recombinant IGF1, I3769, Sigma-Aldrich, Germany). During differentiation, myotubes were cultured in the absence or presence of 20nM, 100nM or 200nM testosterone, 1nM, 5nM or 10nM β-estradiol, 20nM, 100nM or 200nM progesterone (Sigma-Aldrich, Germany) or solvent controls (0.012% ethanol). As there was no difference found between non-treated controls and solvent controls, only non-treated controls are depicted in the figures. Medium was changed three times per week, with or without additional hormones and controls, and 48 hours before harvest. Mycoplasma-free culture conditions and cells are subjected to regular monitoring and analysis using the MycoAlert Mycoplasma Detection Kit (Lonza, Switzerland).

### Quantitative PCR

RNA was extracted employing the RNeasy kit (Qiagen, Germany). For mRNA detection, reverse transcription was carried out with the Transcriptor First Strand Synthesis Kit (Roche, Switzerland). Expression was measured using QuantiFast SYBR Green PCR Mix and QuantiTect Primer Assays (*ATRNL1*: QT00077147; *DDX3Y*: QT01011465; *EIF1AY*: QT01674575; *EIF1AX*: QT00233492; *ENO3*: QT01666434; *GGT7*: QT00058002; *GPD1*: QT00098322; *GRB10*: QT00031276; *IRX3*: QT00227934; *KDM5C*: QT01666931; *LDHA*: QT00001687; *LDHB*: QT00071512; *MYBPC1*: QT00078407; *MYBPC2*: QT00014882; *MYBPH*: QT00000588; *MYH1*: QT01671005; *MYH2*: QT00082495; *MYH3*: QT00068439; *MYH4*: QT01668779; *MYH6*: QT00030807; *MYH7*: QT00000602; *MYL3*: QT00090223; *PLIN2*: QT00001911; *RPS28*: QT02310203; *RPS4Y1*: QT01670613; *TBP*: QT00000721; Qiagen, Germany) in a LightCycler 480 (Roche, Switzerland). Standards were generated by purifying the PCR-product (MinE-lute PCR Purification Kit, Qiagen, Germany) and 10-fold serial dilution. Relative quantification was performed by normalizing expression to the mean of housekeeping genes RPS28 and TBP.

### Statistical analyses

Linear regression models were analyzed with R4.1.1/RStudio ^42^. Normality was tested by Shapiro-Wilk-test from the R package ‘stats’ (v3.6.3) and non-normal data were log-transformed. Differences between groups or time points were calculated using (paired) limma analysis. In the case of p-value adjustment a linear model was calculated (Benjamini-Hochberg post hoc adjustment). For time points comparisons the fold change was calculated. Differences between individual groups were assessed using one-way ANOVA with Fisheŕs LSD post hoc test or Bonferroni correction for multiple comparisons when appropriate. Graphs were made using the R packages ‘ggplot2’ (v3.3.2) and ‘ggrepel’ (v0.9.1) and figures assembled using InkScape (v1.0). Functional enrichment analysis was carried out online using https://biit.cs.ut.ee/gprofiler/ with p<0.05 as threshold and g:SCS as method for multiple testing correction. Homo sapiens was chosen as organism and analysis was performed using the databases from gene ontology biological process (GO:BP), cellular components (GO:CC), molecular functions (GO:MF), Reactome (REAC), KEGG, Wikipathways (WP) and human protein atlas (HPA). MitoCarta 3.0 was used for the annotation to mitochondria.

## Results

### Study cohort characteristics

The total subject group and training regime has been described elsewhere ^36^. As the initial study design was not focused on sex-specific differences our study cohort included more females than males (Table 1). The age range of the subjects was similar for both females and males, with a slight statistical difference. The average age of females was 28 years, while that of males was 33 years. Because of this we adjusted our statistical analysis for age. By study design, all participants had a BMI > 27 (11 participants > 30) with no difference between sexes (Table 1). All had a sedentary lifestyle (<120 min of physical activity per week). At baseline, females and males differed in body fat distribution, cardiorespiratory fitness, exercise performance (Table 1), and estradiol and testosterone serum concentration (Table 2). Females had higher total adipose tissue, subcutaneous adipose tissue, and adipose tissue content in lower extremities, while males had higher visceral adipose tissue and non-adipose tissue content in the lower extremities. Males had higher VO_2_peak/body weight and performance in Watt [W] (Table 1). The response in physical fitness and body fat distribution to the 8-week training intervention was highly comparable between sexes, with an increase in the individual aerobic threshold and VO_2_peak and a reduction in adipose tissue compartments. The only statistically significant differences between females and males are found in the fold changes in adipose and non-adipose tissue of the lower extremities. Considering the individual changes over 8 weeks, only the increase in non-adipose tissue of females was statistically significant.

**Table 2.** Sex hormones.

|  | Baseline |  |  | Trained |  |  | Fold change |  |  |
| --- | --- | --- | --- | --- | --- | --- | --- | --- | --- |
|  | Female | Male | p-Value | Female | Male | p-Value | Female | Male | p-Value |
| **Estradiol** | 342 | 162 | **0.004** | 393 | 202 | **0.008** | 1.37 | 1.24 | 0.647 |
| **[pmol/L]** | ± 213 | ± 25.1 |  | ± 225 | ± 81.5 |  | ± 0.87 | ± 0.46 |  |
| **Progesterone** | 3.46 | 1.81 | 0.111 | 7.02 | 1.82 | 0.063 | 5.21 | 1.06 | 0.127 |
| **[nmol/L]** | ± 3.79 | ± 0.76 |  | ± 9.97 | ± 0.58 |  | ± 9.91 | ± 1.06 |  |
| **Testosterone** | 1.59 | 13.3 | **<0.001** | 1.61 | 14.2 | **<0.001** | 1.02 | 0.23 | 0.712 |
| **[nmol/L]** | ± 0.50 | ± 4.04 |  | ± 0.63 | ± 5.07 |  | ± 0.29 | ± 0.29 |  |

### Pronounced differences on multi-OMICs level between female and male skeletal muscle at baseline

We analyzed the skeletal muscle transcriptome, proteome and methylome data obtained from the skeletal muscle biopsies collected in the resting state before the 8-weeks endurance exercise intervention for sex differences. Not considering sex-specific differences, the methylome and transcriptome data from this cohort were reported recently ^44^. We now identified 100,515 differentially methylated CpG sites in 16,012 genes (p< 0.05) and 1,366 differentially abundant transcripts (p<0.05) in female and male skeletal muscle (Figure 1 A). The proteome data set comprised 1,857 proteins identified with high confidence and quantified in at least 90% of all samples at one time point. We identified 120 differentially abundant proteins (p<0.05) between female and male skeletal muscle at baseline (Figure 1 A). Out of all differentially methylated CpG sites, 84 240 (83.8%) CpG sites were hypermethylated in females when compared to males and only 16,275 (16.2%) hypomethylated. DNA methylation in promoter and enhancer regions is associated with silencing of gene transcription, whereas high DNA methylation levels are typically found in the body of actively transcribed genes ^48^. Considering the genomic position of the DNA methylation site and the direction of change in methylation, differential expression of 802 (58.7%) transcripts could be linked to differences in DNA methylation. The high proportion of elevated methylation levels in female participants was similarly detected in the promoters and gene bodies of the corresponding 802 genes (Figure 1 B). Sex-specific abundance of 80 (66.6%) proteins could be linked to differences in DNA methylation, and a total of 19 differentially abundant proteins could be traced back over differential gene expression to sex-specific DNA methylation (Figure 1 A). Our data suggest that distinct epigenetic signatures between females and males could potentially explain up to 60% of the sex-dependent changes in skeletal muscle transcriptome/proteome at baseline. Thus, deeply rooted differences exist in untrained skeletal muscle of sedentary, overweight/obese females and males spanning from the proteomic over transcriptomic down to the epigenomic level.

**Figure 1.**
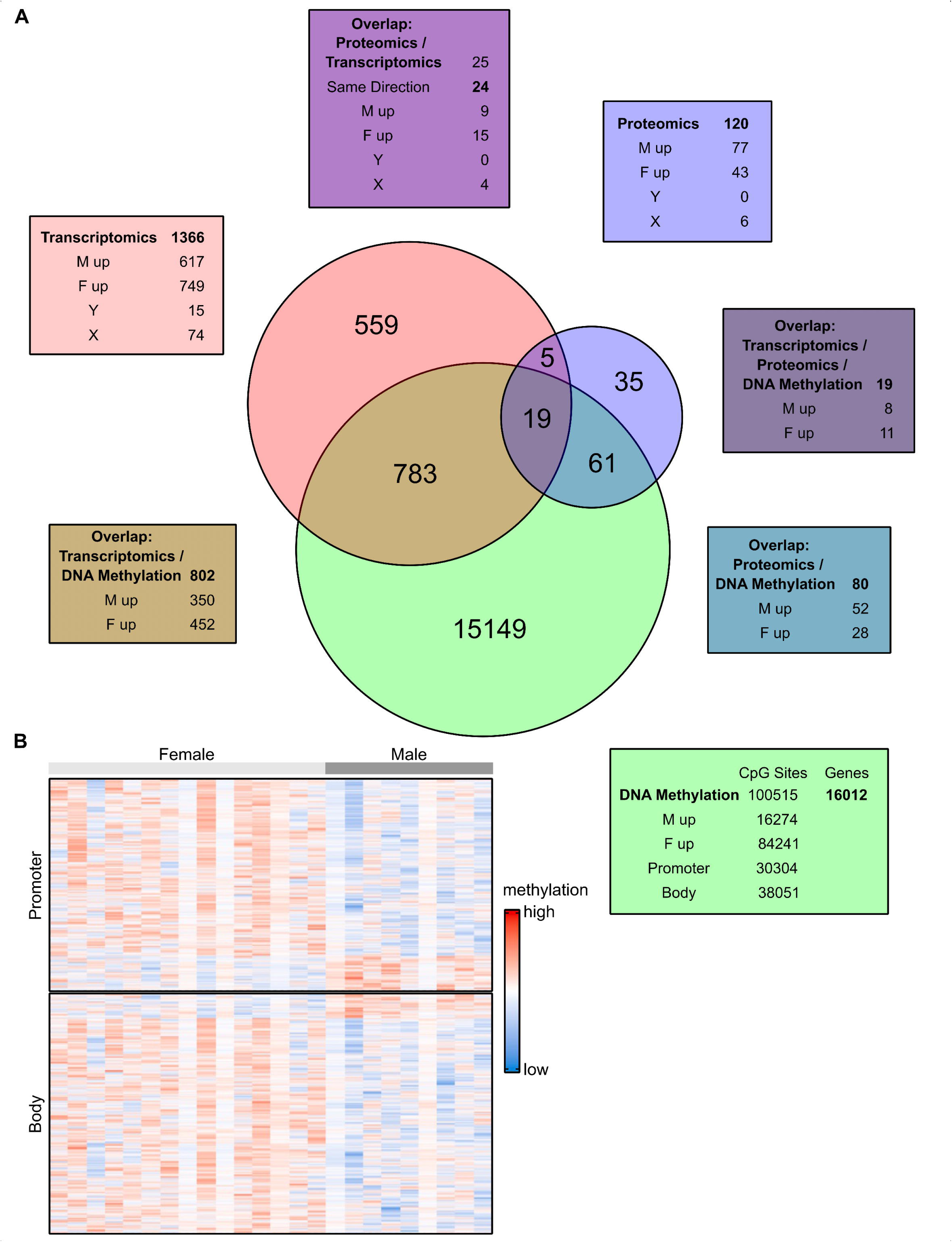
Multi-Omics analysis of female vs. male skeletal muscle at baseline A) Skeletal muscle biopsies obtained at baseline were analyzed for transcriptomic, proteomic and epigenomic (DNA-methylation) differences between females (F) and males (M). Numbers show the sex differences found in the datasets as indicated. Venn diagram visualizing the overlap of the differences in the multi-omics datasets. B) Heatmap depicting scaled methylation levels of 2,192 CpG sites located in 802 differentially expressed genes. Each column represents a skeletal muscle sample from an individual donor, each row indicates the methylation level of a single CpG site with significant differences between males and females. Rows marked in light grey represent female participants, male participants are indicated in dark grey. The differentially methylated CpG sites found in the promoters are displayed in the upper part of the heatmap, those located in gene bodies are shown in the lower part. Statistical significance was determined by limma t-test p < 0.05, n=25 (16f/9m).

### Transcriptomic differences point to an influence of sex on glucose and lipid metabolism in skeletal muscle

In the transcriptome, the most pronounced differences were found in sex chromosomal transcripts dominated by ChrY-located transcripts in males (Figure 2 A). In contrast, higher expression of ChrX-located transcripts was evenly distributed among sexes. The majority of transcriptomic differences was found in autosomal genes (Figure 1, Figure 2 A). The transcriptomic differences were validated by comparison with the skeletal muscle transcriptome data of a previously published independent second cohort consisting also of sedentary subjects with overweight/obesity participating in an exercise intervention study ^49^. The overlap yielded 196 transcripts differentially expressed between sexes and confirmed the differential expression pattern of sex chromosomal encoded transcripts (Figure 2B). Among top candidates of differentially expressed transcripts conserved in both cohorts were genes with key functions in glucose and lipid metabolism such as *GRB10*, *LDHB*, and *LPL* (higher in females) and *ALDH1A1*, *PFKFB1, PGK1*, and *PHKA1* (higher in males) (Figure 2 B,C). Enrichment analysis of the sex-specific transcripts differentially expressed in skeletal muscle between females and males pointed to differences in glycolysis and demethylase activities (Figure 2 D).

**Figure 2.**
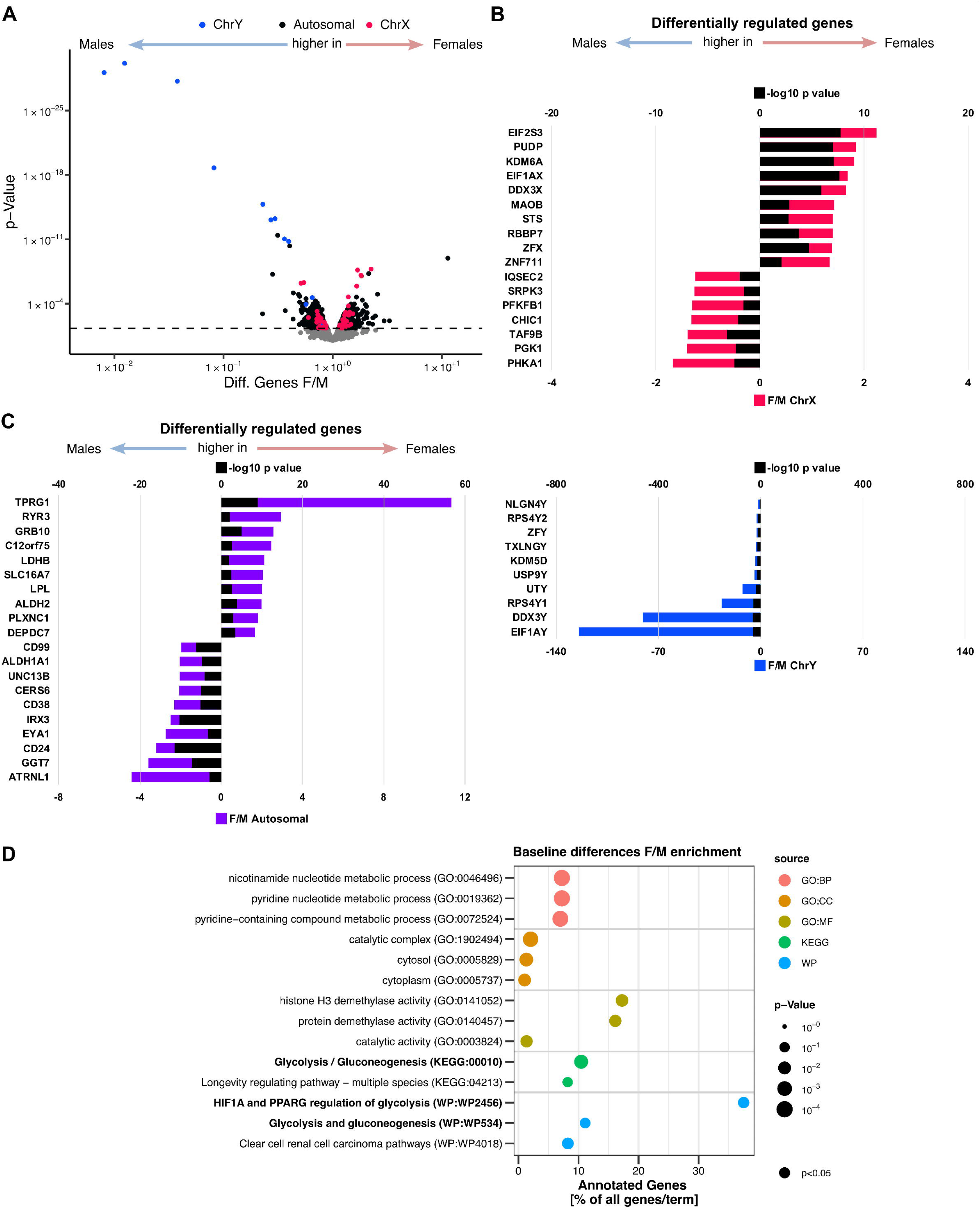
Transcriptomic analysis of female vs. male skeletal muscle at baseline Skeletal muscle biopsies obtained at baseline were analyzed for transcriptomic differences between females and males. A) Volcano plot of differentially expressed transcripts being higher in males (left) or higher in females (right), X-chromosomal genes are labeled in red, Y-chromosomal genes are labeled in blue. Top 10 transcripts higher expressed in males (left) and females (right) conserved in this and another independent cohort ^49^are plotted, fold change (F/M) in B) X-chromosomal (red) and Y-chromosomal (blue) genes, C) autosomal (purple) (bottom axis),-log10 p-values are plotted in black (top axis). D) Enrichment analysis based on the 196 differentially expressed genes in both cohorts. Statistical significance was determined by limma t-test p < 0.05, n=25 (16f/9m) and validation cohort (n=19; 12f/6m).

### Proteomic differences provide a molecular basis for sex-specificity of glucose and lipid utilization and fiber type prevalence

In the skeletal muscle proteome, enrichment analysis of the 120 proteins differentially abundant between females and males revealed that these proteins were mainly associated with striated muscle contraction and metabolic pathways, in particular glucose metabolism (Figure 3 A). While all lipid metabolism-associated proteins (ACSL1, CD36, PLIN2, PLIN4) were more abundant in female muscle, all glycogen degradation and glycolysis-associated enzymes were more abundant in skeletal muscle of males (Figure 3 B). Among these are many key enzymes of glycogen degradation and glycolysis, and differential expression was also found on transcriptomic level (Figure 3 C). Proteins with mitochondrial localization, based on MitoCarta 3.0, exhibited an even distribution between sexes, with no clear tendency for either males or females to have a higher abundance of these proteins (Figure 3 D). The clear indication of elevated glucose/glycogen utilization in male muscle that can be associated with the fast-twitch 2A and 2X muscle fibers raised the question on the distribution of differentially abundant proteins that can be specifically attributed to slow and fast-twitch fibers. Thus, we overlapped our proteomic results with the list of human fiber type-specific proteins reported by Murgia et. al. in 2021 ^31^ (Figure 3 E). Indeed, almost all differentially abundant proteins with preferential expression in type 2 fibers were more abundant in male skeletal muscle including many of the enzymes of glycogen degradation and glycolysis while proteins specific for type 1 oxidative fibers were more abundant in female skeletal muscle (Figure 3 E). To conclude, the proteome data obtained from the skeletal muscle biopsies before the intervention provide a strong molecular basis for the reported higher percentage of type 2 fibers in male muscle with a preference for carbohydrate utilization and the higher percentage of type 1 fibers in female muscle supporting a preference for lipid utilization. Of the proteins with fiber type-specific preference (Figure 3 E) only proteins with metabolic function (ENO3, GPD1, LDHA, PGK1, TPI, ACSL1, CD36) and calsequestrin CASQ2 were also different at transcript level (Supplementary Data Table 1), while the sex-specific abundance of proteins representing the type 1 and 2 fiber-specific contractile and structural profile appears to be post-transcriptionally regulated.

**Figure 3.**
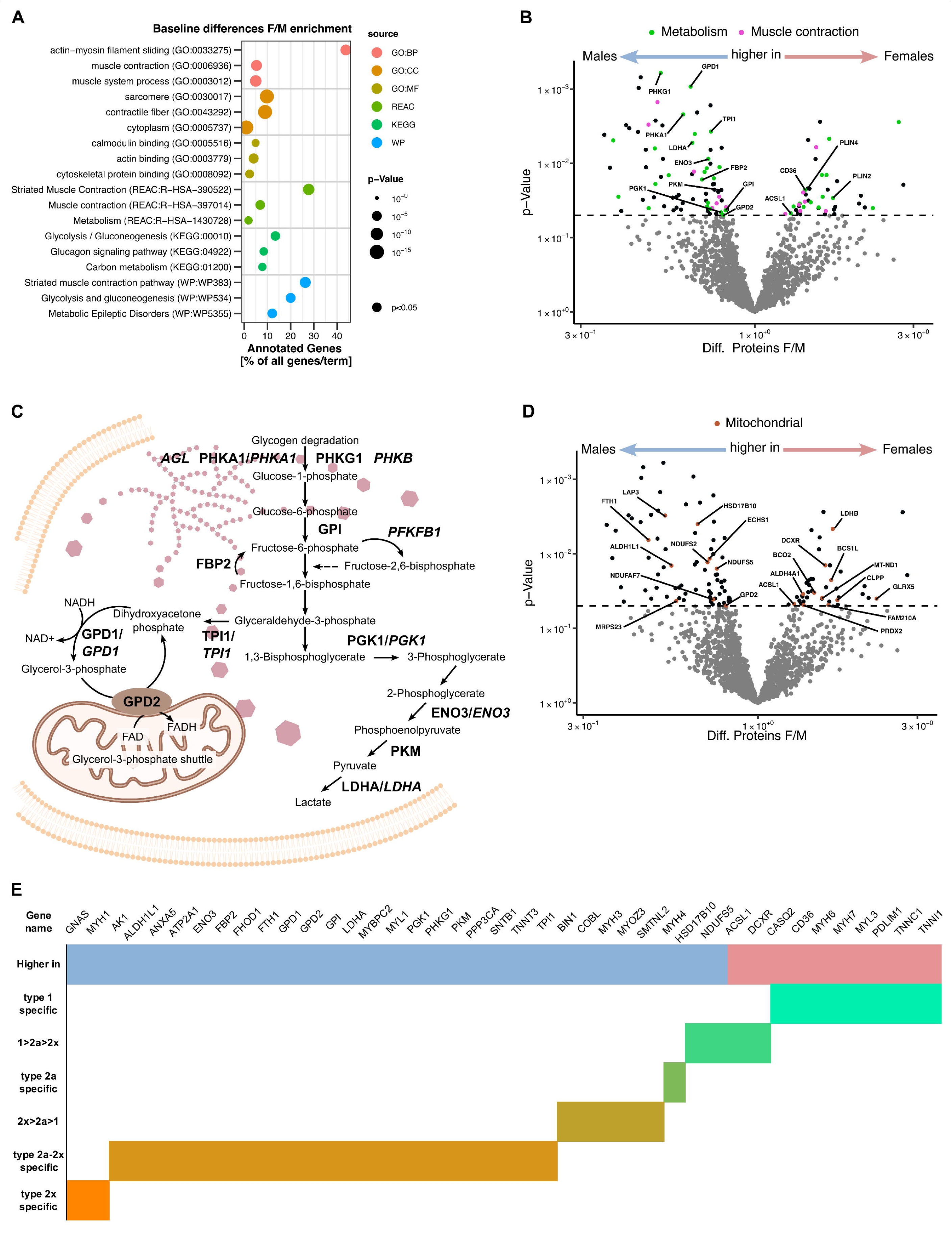
Proteomic analysis of female vs. male skeletal muscle at baseline Skeletal muscle biopsies obtained at baseline were analyzed for proteomic differences between females and males. A) Enrichment analysis based on the 120 differentially abundant proteins between females and males. B) Volcano plot of differentially abundant proteins being higher in males (left) or higher in females (right). Color-coded for proteins enriched in the top terms (REAC database) Metabolism (green) and Muscle contraction (pink). Key enzymes of glycogen, glucose and lipid metabolism are labeled. C) Schematic representation of the glycogen degradation and glycolysis pathway and the key enzymes (PROTEIN/TRANSCRIPT) found to be elevated in male skeletal muscle. D) Volcano plot of differentially abundant proteins being higher in males (left) or higher in females (right) as shown in (A), proteins with mitochondrial localization are labeled. E) Table of proteins with differential abundance between females and males (red higher in females, blue higher in males), with higher percentage of abundance in oxidative slow type 1 fibers and fast type 2a and 2x fibers based on ^31^.Statistical significance was determined by limma t-test p < 0.05, n=25 (16f/9m).

### Hormonal regulation of sex-specific differences observed in vivo in myotubes in vitro

To analyze whether the observed differences in skeletal muscle between females and males at baseline are conserved in myotubes obtained from female and male donors, we cultivated satellite cells isolated from the baseline muscle biopsies of the study participants and differentiated the myoblasts for 7 days into fully differentiated myotubes and analyzed the mRNA expression of top differentially expressed genes (Figure S 2 A). Expression of genes located on the X chromosome like *EIF1AX* and *KDM5C*, higher in female skeletal muscle, was conserved in the cultured myotubes (Figure S 2 B). For autosomal genes we analyzed candidates out of the top 10 transcripts shown in Figure 2B and of the metabolically relevant transcripts shown in Figure 3B. For transcripts higher in the female muscle represented by *LDHB* and *GRB10*, a slight trend for higher abundance in female myotubes was only observed for *GRB10* (p=0.088) (Figure S 2 C). Genes exhibiting a higher expression in male skeletal muscle like *ENO3*, *LDHA*, *GPD1, ATRNL1*, *GGT7* and *IRX3* were not higher expressed in the respective male myotubes (Figure S 2 D).

As transcriptomic differences between cultured female and male myotubes were limited to non-autosomal genes we investigated whether the differences found in vivo were inducible by the presence of the sex hormones estradiol, progesterone or testosterone during myotube differentiation (Figure 4 A). Myoblasts were treated for 7 days with hormone concentrations that were based on the physiological serum concentrations of either females or males or 5 to 10 times higher. In our analysis we focused on muscle fiber type-specific and metabolically relevant genes. The slow-twitch oxidative fiber type-specific protein MYH7 was slightly induced by 200 nM progesterone on transcriptional level, and *MYL3* and *MYH6* transcripts were reduced by 20, 100, and 200 nM testosterone reflecting the lower protein abundance in male muscle (Figure 4 B-G). The transcripts of fast-twitch fiber type-specific proteins MYBPH and MYBPC2 were increased by 20, 100 and 200 nM testosterone reflecting the higher protein abundance in male muscle (Figure 4 H, I, L, M). A strong inducing effect of testosterone was found on *MYBPC1,* the MYBPC1 protein however, is considered to be more abundant in slow-twitch fibers and was not different in female and male skeletal muscle biopsies (Figure 4 J, K). No effect of either hormonal treatment was found on *MYH1, MYH*3 and *MYH4* transcripts, which are considered as fast-twitch fiber type-specific proteins (Figure S 3 A, C, D). Transcript levels of fast-twitch fiber type MYH2 were even reduced under testosterone treatment (Figure S 3 B). Many other differences observed in skeletal muscle of females and males could also not be evoked as expression of *LDHA*, *LDHB*, *IRX3*, *GPD1*, *ENO3* and *PLIN2* were not regulated by sex hormone treatment, except a reduced expression of *LDHA* at 10 nM estradiol (Figure S 3 E-J). None of the analyzed transcripts was expressed in a sex-specific manner in untreated myotubes obtained from female or male donors. Among the sex hormones tested, only testosterone induced transcriptomic changes at a concentration found in serum of males. In conclusion, there is little evidence for in vitro conservation. The treatment of human myotubes with sex hormones can, to some extent, restore the in vivo sex differences.

**Figure 4.**
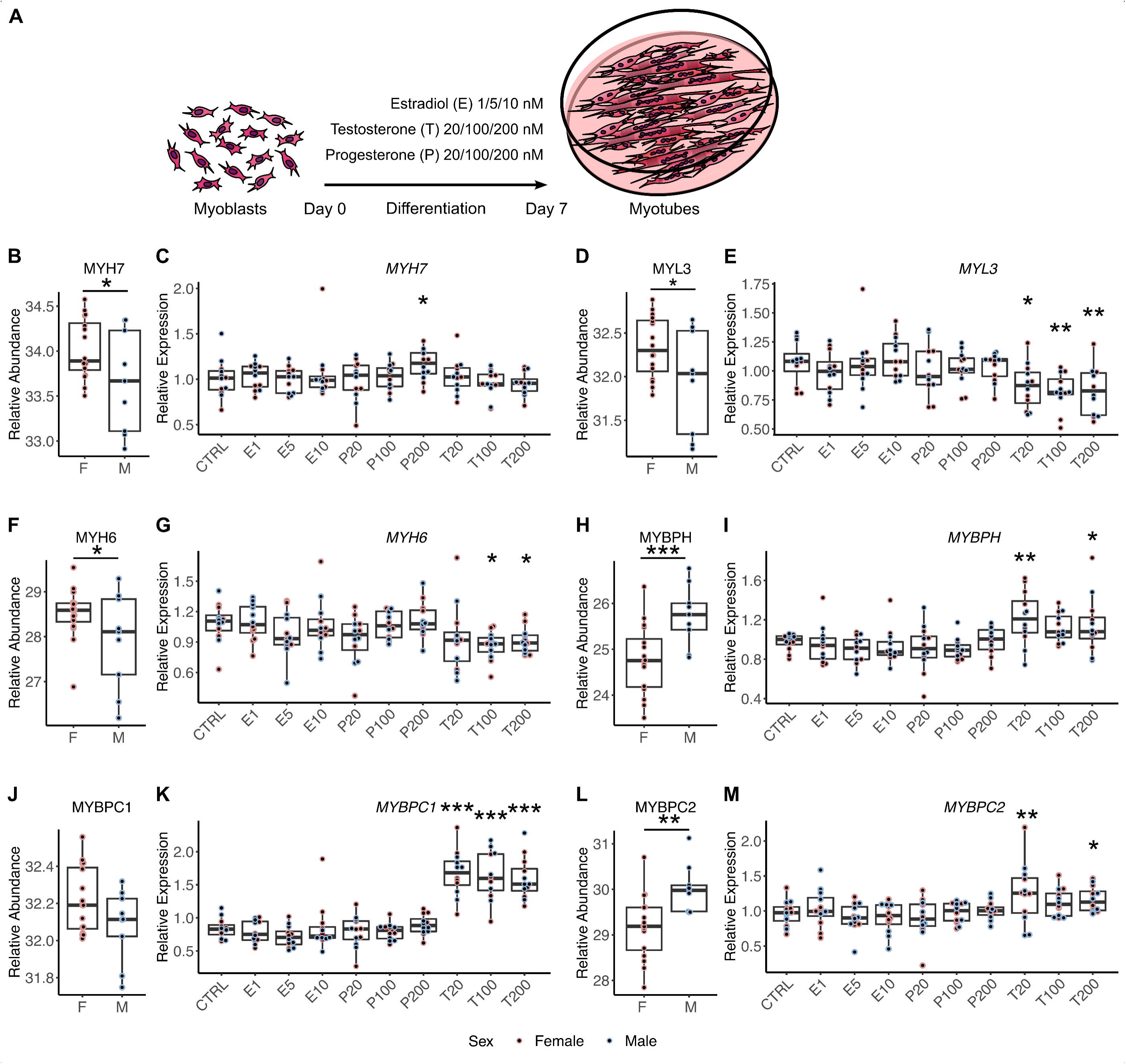
Sex hormone-driven transcriptional regulation of fiber type-specific proteins in myotubes in vitro A) Myoblasts derived from 6 female and 6 male donors were cultured and differentiated into myotubes for 7 days. During differentiation, myotubes were treated with estradiol (E) (1/5/10nM), progesterone (P) (20/100/200nM) or testosterone (T) (20/100/200nM) or left untreated (CTRL). RNA expression of C) MYH7, E) MYL3, G) MYH6, I) MYBPH, K) MYBPC1 and M) MYBPC2 in myotubes after hormonal treatment, respectively. Differences in RNA expression were determined by one-way ANOVA BonferronI, n=12 (6f/6m), * p<0.05, ** p<0.01, *** p<0.001. To the left is the respective protein abundance in skeletal muscle of females (F) and males (M) at baseline (B, D, F, H, J, L). Statistical significance was determined by limma t- test p < 0.05, n=25 (16f/9m), * p<0.05, ** p<0.01, *** p<0.001. Red dots, female donor; blue dots, male donor.

### Sex influences the transcriptional response to acute exercise

In light of the deeply rooted differences in skeletal muscle of females and males at baseline, we expected that the initial response of skeletal muscle to acute exercise differs between sexes as well. We focused on the transcriptomic response since the skeletal muscle biopsies were collected 60 min after the first 30 min-ergometer exercise bout, when particularly a transcriptional regulation response was observed (Figure S 4 A). The comparison of the transcriptional response in female and male skeletal revealed that females upregulated 274 transcripts in response to the first acute exercise bout which were not regulated in males. The transcripts were enriched in pathways of glucose metabolism, glycolysis, pyruvate metabolism and TCA cycle (Figure 5 A). Male skeletal muscle showed a distinct transcriptomic response. The upregulated 87 transcripts, which were not regulated in females, were enriched in mitochondrial and oxidative stress-related pathways (Figure 5 B). This includes pronounced upregulation of cellular stress-inducible transcription factors *ATF3* and *JUN,* protein kinase *STK39*, the divalent metal transporter *SLC39A14*, and oxidative stress-responsive *HMOX*, *MT1A* and *MT1B* transcripts (Figure 5 C-I). Serum myoglobin concentrations serve as marker of increased myofiber damage and skeletal muscle membrane vulnerability. In line with elevated transcriptional markers of cellular stress in male skeletal muscle, serum myoglobin levels were increased after the acute exercise bout only in males (Figure 5 J). Baseline myoglobin values were also by trend higher in males than in females (p=0.08), presumably due to the higher muscle mass of males. Plasma lactate was increased to a similar concentration in both sexes (Figure 5 K).

**Figure 5.**
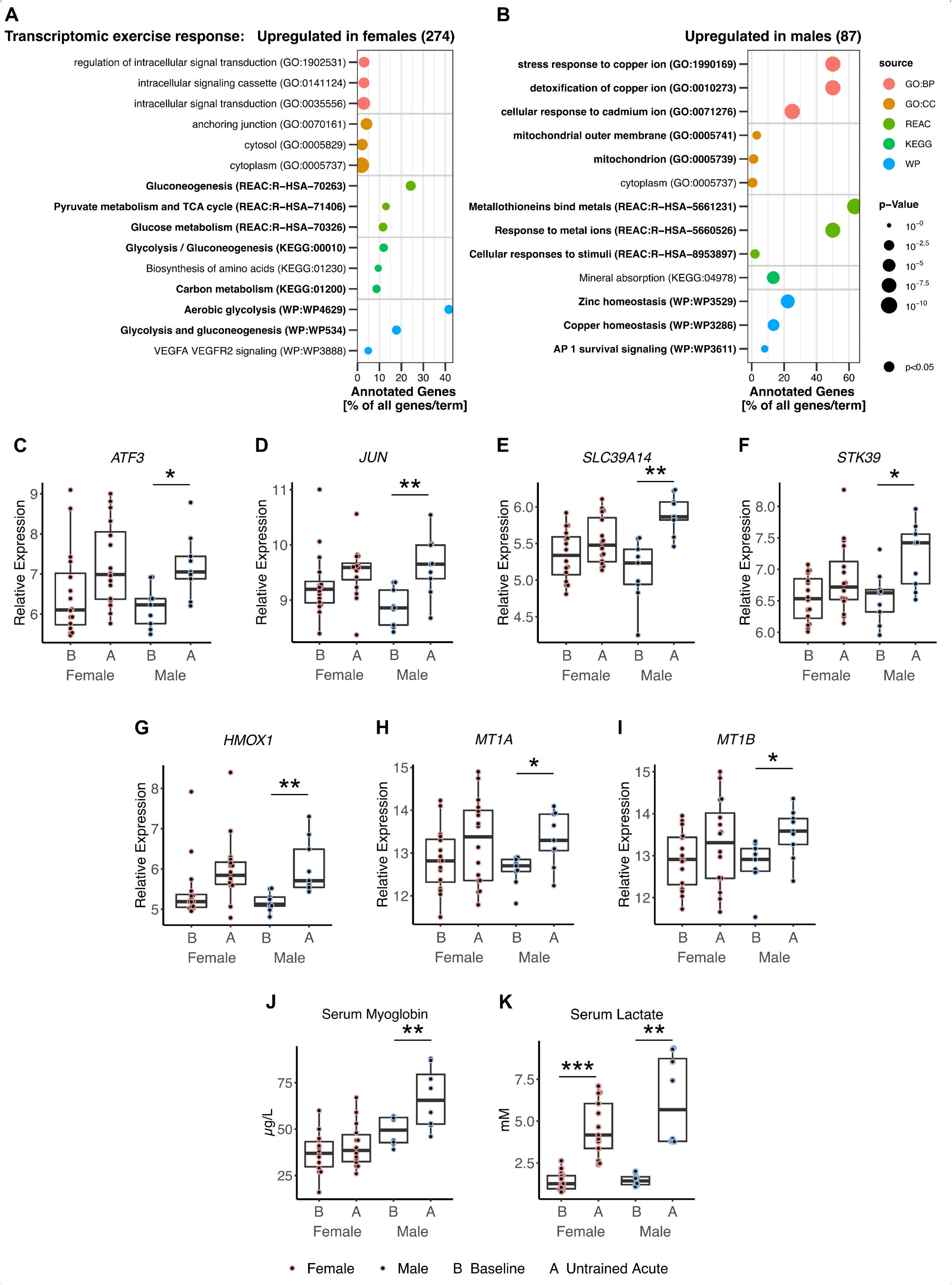
Differences in the transcriptional response of female and male skeletal muscle to acute exercise Enrichment analysis based on A) 274 transcripts upregulated in female skeletal muscle only and B) 87 transcripts upregulated in male skeletal muscle only in response to acute exercise. Relative transcript abundance of C) ATF3, D) JUN, E) SLC39A14, F) STK39, G) HMOX1, H) MT1A and I) MT1B in skeletal muscle of females and males at baseline (B) and 60min after the first acute exercise bout (A). Statistical significance was determined by limma t-test BH, p < 0.05, n=25 (16f/9m), * p<0.05, ** p<0.01. Serum myoglobin J) and lactate K) was measured in females and males at baseline (B) and 5min after the first acute exercise bout (A). Statistical significance was determined by one-way ANOVA LSD, n=22 (14f/8m), ** p<0.01.

The acute exercise bouts were performed at an individual intensity corresponding to 80% of VO_2_peak. Accordingly, heart frequency per minute (159±12 vs. 150±17, p=0.188) and rating of perceived exertion of leg work (15 vs. 16, p=0.518, corresponding to BORG scala) during the first 30 min-ergometer exercise bout were not different between sexes, despite the lower absolute intensity in females (101±20 vs. 143±20 W, p<0.001). Of note, the absolute intensity of the performed exercise bout did not correlate with the upregulation of *ATF3*, *JUN*, *STK39*, *SLC39A14*, *HMOX*, *MT1A* and *MT1B* transcripts (Supplementary Table 1).

### Eight weeks of endurance training increase key enzymes of mitochondrial metabolism in both sexes and equalize baseline differences

To elucidate the sex-specific response to the 8-week endurance training intervention, we focused on the proteomic response. As the skeletal muscle biopsies were obtained 5 days after the final training session, mainly sustained changes in the proteome, but fewer changes in the transcriptome were detected (Figure S 4 A). Unlike to the observed differences in skeletal muscle of females and males in response to the first acute exercise bout, after 8 weeks of supervised endurance training, the proteomic response was highly comparable in both sexes. Both females and males upregulated 185 proteins enriched in oxidative phosphorylation, mitochondrial respiration and ATP synthesis (Figure 6 A). Common protein responses to the 8-weeks training included upregulation of mitochondrial marker proteins and key enzymes of the TCA cycle, respiratory chain and β-oxidation as exemplarily shown for CS, MT-CO1, MT-CYB and CPT1B (Figure 6 B-E). Among sex-specific responses were alterations in key enzymes of acetyl-CoA metabolism. Cytosolic acyl-CoA synthetase short chain ACSS2, which is involved in lipid synthesis, was reduced only in female muscle, whereas the mitochondrial ACSS1 was increased only in male muscle after 8 weeks of training (Figure 7 F, G). The key enzyme of ketogenesis HMGCS2 was upregulated only in female muscle (Figure 6 H). Among the proteins found to be reduced only in male muscle in response to the 8-weeks training were the fast-twitch and glycolytic fiber type-specific proteins ALDH1L1, MYBPH, MYH1 and MYH3, resulting in equalized protein levels between sexes after training (Figure 6 I-L). Based on this observation, we reevaluated all fiber type-specific proteins that were initially different between females and males at baseline (Figure 3 D) and found that most of those were no longer differentially abundant after 8 weeks of endurance exercise (Figure 6 M). Similarly, the abundance of enzymes involved in glycogen degradation and glycolysis that were initially higher in males at baseline (Figure 3 B) was reduced in male muscle toward the levels of females (Figure 6 M). In summary, 8 weeks of controlled endurance exercise equalized initially observed differences in skeletal muscle toward a common metabolically beneficial response in females and males.

**Figure 6.**
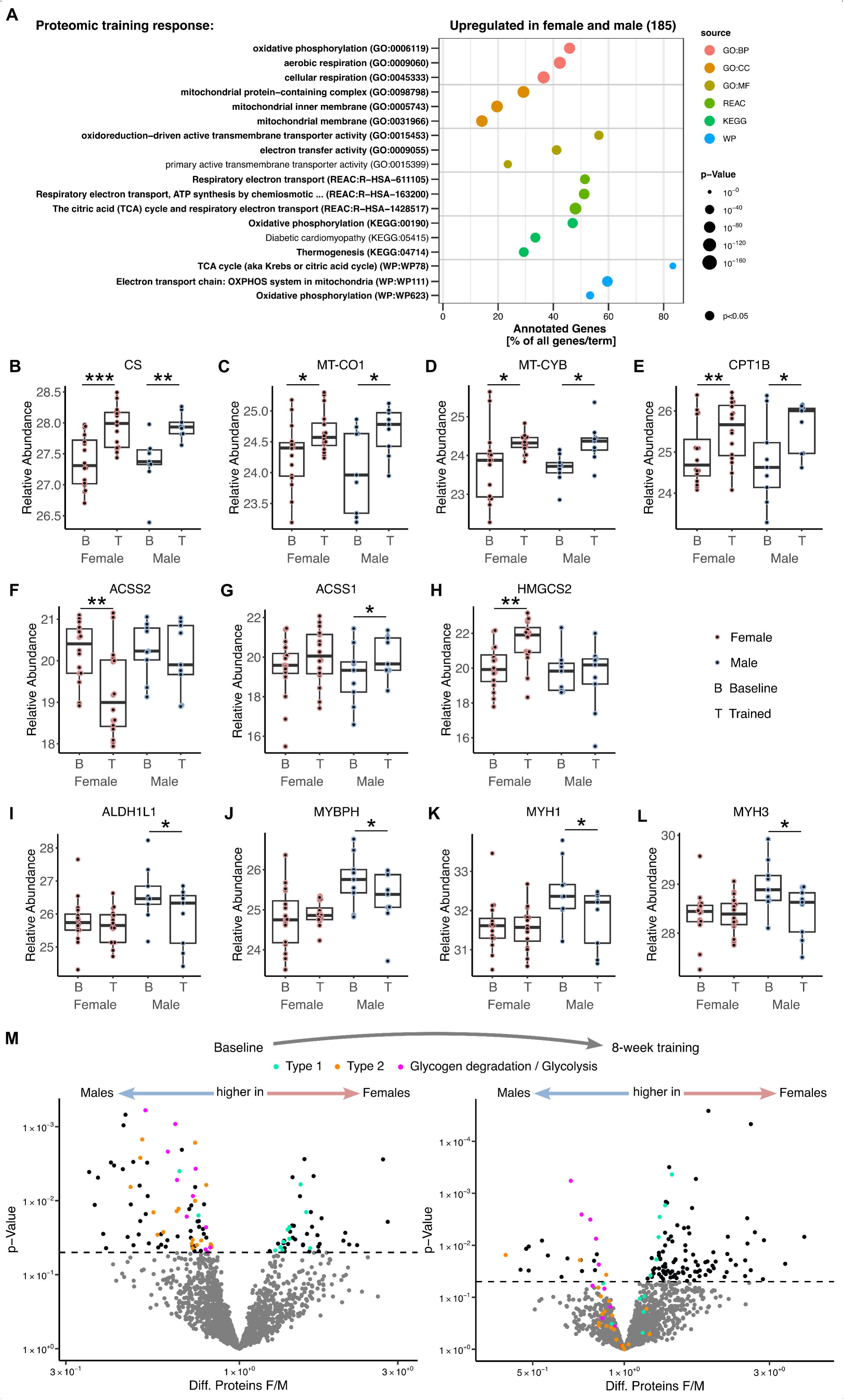
Effects of 8-week training on skeletal muscle proteome of females and males A) Enrichment analysis based on 185 proteins upregulated in both female and male skeletal muscle in response to 8 weeks of training. Relative protein abundance of B) CS, C) MT-CO1, D) MT-CYB, E) CPT1B, F) ACSS2, G) ACSS1, H) HMGCS2, I) ALDH1L1, J) MYBPH, K) MYH1 and L) MYH3 in skeletal muscle of females and males at baseline (B) and after 8 weeks of training (T). K) Volcano plots of differentially abundant proteins at baseline and after 8 weeks of training, being higher in males (left) or higher in females (right), color-coded for differentially abundant proteins at baseline, turquoise for oxidative slow type 1 fibers and orange for fast type 2 fibers based on (Murgia et al. 2021). Proteins relevant for glycogen degradation / glycolysis were shown as pink dots. Statistical significance was determined by limma t-test, p < 0.05, n=25 (16f/9m), * p<0.05, ** p<0.01, *** p<0.001.

## Discussion

In this study, we provide for the first time a yet missing comprehensive picture of molecular differences between female and male skeletal muscle by applying a multi-omics approach employing DNA methylation, transcriptomics and proteomics. We investigated not only the differences in untrained muscles, but also the impact of sex on the response to acute exercise and to an 8-week endurance training by analyzing three skeletal muscle biopsies per subject obtained before training, after the first exercise bout and 5 days after the final exercise session.

A few studies have focused on analyzing sex-specific DNA methylation in skeletal muscle, including the X chromosomes ^50,51^, while others have concentrated solely on CpG sites located in autosomal chromosomes ^33^. In both strategies of methylome analysis, just like in our skeletal muscle data, overall higher methylation levels were detected in females. To examine genomic regions with consistent and extended differential methylation, Landen et al. investigated differentially methylated regions (DMRs) in the skeletal muscle of 222 males and 147 females ^33^. They identified 10,240 DMRs, with 94% hypermethylated in females. Davegardh et al. reported not only hypermethylation of DMRs in skeletal muscle biopsies of females, they also found a higher methylation of 62% of DMRs in autosomal genes in human myoblasts obtained from female donors. We showed that the hypermethylation of DMRs is evenly distributed between the promoter region and the body of genes with a transcriptional regulation that aligns to the differences in DNA methylation. Thus, the hypermethylation of CpG sites in female muscle appears to be a consistent finding and is even conserved in myoblasts. The extend of hypermethylation in female skeletal which is independent from the genomic location suggest a global difference in enzymatic methylation or demethylation activities, but the reason for this sexual dimorphism is unknown. Landen et al. could not validate a relation of circulating sex hormone concentrations and the hypermethylation in females. The observation is not restricted to skeletal muscle, hypermethylation of autosomal genes was also reported in blood leukocytes of females and umbilical cord blood of newborn girls ^52,53^. The overlap of the transcriptome and methylome data at baseline revealed that close to 60% of differentially expressed genes and proteins between females and males are potentially mediated by sex-specific DNA methylation patterns. Thus, our data suggest DNA methylation as one potential mechanism, among others, that contributes to metabolic differences between females and males in skeletal muscle prior exercise intervention.

Sexual dimorphisms in the transcriptome of skeletal muscle have been also investigated by others ^32–34^. The studies varied in health status and age of the participants and used either microarray data as we did ^33,34^ or RNA sequencing data ^32^. Comparison of our 196 differentially expressed transcripts in the two exercise cohorts with the most recent sequencing data of the young group (18-30 years) in the study of Pataky et al. ^32^ revealed an overlap of 141 transcripts among them 111 autosomal genes. The sex-dependent regulation of lipoprotein lipase *LPL*, acyl-CoA synthetase short chain *ACSS2*, insulin receptor *INSR*, growth factor receptor-bound protein *GRB10*, the key antioxidant enzyme catalase *CAT* and ryanodine receptor type 3 *RYR3* was not only reported by Pataky et al. but also found in a lean cohort by Welle et al. ^34^. The higher abundance of *LPL* and *ACSS2* in females reflect the increased capacity for triglyceride clearance from lipoproteins and fatty acid synthesis in female muscle. GRB10 is a modulator of the proximal IGF1 and insulin signaling, and its ablation in skeletal muscle leads to muscle hypertrophy in mice and evidence for reduced insulin signaling ^54^. A higher GRB10 expression in females could potentially contribute to the smaller muscle mass in females by dampening muscle growth signals. On the contrary, higher insulin sensitivity was reported for female skeletal muscle ^55^, which would be in line with higher abundance of the insulin receptor. A higher enzyme activity of catalase was found after estradiol replacement therapy in skeletal muscle of rats ^56^ and can be one factor responsible for the stress resilience observed in female skeletal muscle after acute exercise. In male muscle, common upregulation is seen for the glycolytic enzymes *ENO3*, *GPD1*, *LDHA* and *TPI1*, again highlighting the sexual dimorphism in glucose metabolism. Highly conserved is also the upregulation of *IRX3* and *CERS6* in males, which is also in line with a differential DNA methylation of the genes between sexes ^32–34^. CERS6 belongs to the ceramide synthases and its higher abundance in male muscle can be implicated in the reported higher susceptibility of male skeletal muscle to develop lipid-induced insulin resistance ^55,57^. One of the top differentially expressed transcripts in this study and others, is *IRX3* (Iroquois Homeobox Protein 3). We found a 2.5-fold higher transcript level in male skeletal muscle alongside a hypomethylated site in the promoter and a hypermethylated site in the gene body in male participants. IRX3 showed also sexual dimorphic expression in skeletal muscle of mice with higher levels in males ^58^. While this report discussed IRX3 expression as potentially androgen-regulated, our data did not show upregulation of IRX3 transcripts in testosterone-treated myotubes irrespective of the sex of the donors. The function of the transcription factor IRX3 was mainly studied in adipose tissue, since the obesity-related genetic variants in the FTO gene affects adipose tissue functions via modulating the expression of IRX3 ^59^. Its function in skeletal muscle is unknown. Thus, it is an interesting question for future research whether IRX3 is involved in the sexual dimorphic morphology and function of skeletal muscle and what causes the differential expression of IRX3 in skeletal muscle of males and females.

Corroborating our findings on the next level of this multi-omics analysis, our proteome data indicate a clear sexual dimorphism in handling energy metabolism in skeletal muscle with a strong preference towards elevated lipid metabolism in females and glycogen degradation and glycolysis in males. Our results are well in accordance to an investigation in non-overweight, untrained females and males where females exhibited higher levels of proteins involved in fatty acid degradation and amino acid metabolism, while males showed more proteins related to carbohydrate metabolism and proteasome pathways ^35^. In addition, our data indicate a sex-specific epigenetic and transcriptional regulation responsible for the elevated levels of enzymes PHKA1, PGK1, TPI1, GPD1 and LDHA in males and elevated PLIN4, CD36 and ACSL1 in females, since the proteomic differences were reflected by differences in the respective transcript levels and in DNA methylation of the genes. This deeply rooted sexual dimorphism in glucose and lipid metabolism is well in accordance with the reported differences in fiber type composition between male and female skeletal muscle ^18,27–30,60^. Our proteome data clearly show a type 1 fiber protein profile in female and a type 2 protein profile in male skeletal muscle and strongly support the results of histological analyses on a proteomic level. The higher proportion of slow-twitch oxidative type 1 fibers in females support endurance exercise and resistance to fatigue. Accordingly, there is some evidence that performance by sex measured as average speed did not differ so much in longer running distance compared to shorter runs ^61^. Males have more fast-twitch type 2 fibers, providing higher power output and faster contraction rates, which can contribute to the greater muscle strength and better sprint distance performance in males which is also due to the difference in muscle mass ^62^. In contrast to the proteome analysis, our transcriptomic data did not hint to a difference in fiber type composition, in line with previous transcriptome studies that also did not highlight fiber type differences between sexes ^32,33^. While the abundance of many fiber type-specific metabolic enzymes and regulators can be attributed to similar differences on a transcriptional and also epigenetic level, this was not observed for the proteins building the contractile apparatus. This raises the question of post-transcriptional mechanisms responsible for the sexual fiber type dimorphism. Among the 120 proteins different at baseline are also proteins involved in translational mechanisms such as translation initiation factors, ribosomal proteins, and a ribosome maturation factor, but a specific contribution of these factors to a divergent translation of sex-specific contractile structure proteins needs to be elucidated.

Many of the described sex-specific differences in skeletal muscle can be controlled by sex hormones. We investigated this further in a donors’ sex matched *in vitro* approach. When looking at several of our top differentially abundant transcripts and proteins in cultured and differentiated myotubes obtained from the participants of the study, we observed conservation of differential expression in transcripts located on sex chromosomes, whereas differences in autosomal located transcripts were not observed in myotubes *in vitro*. A comparison of the genome-wide transcriptomic profile in human myotubes obtained from 13 female and 13 male donors reported 22 genes with sex-specific differential expression among them 13 transcripts of autosomal genes ^50^. None of these 13 transcripts was found in our comparison of the skeletal muscle transcriptomes of females and males, underlying the weak conservation of sex-specific autosomal differences comparing skeletal muscle tissue and myotubes *in vitro*. Accordingly, a difference in glucose and fatty acid handling was not observed in human myotubes obtained from female and male donors, quite in contrast to the *in vivo* data ^63^. One obvious difference between skeletal muscle tissue and cultured skeletal muscle cells is the lack of sex-specific hormonal regulation. Previous studies already investigated human myotubes after sex hormone treatment and reported a shift in metabolism away from glucose storage and glucose oxidation towards lipid oxidation when myotubes were acutely treated with supraphysiological 10µM testosterone or 10µM estradiol ^64,65^. These effects were independent from the sex of the donors. Similarly, when human myotubes were treated for 4 days with 35nM testosterone or 1nM estradiol corresponding to physiological serum concentrations, male and female myotubes exhibited comparable transcriptional responses ^32^. This is in line with our current data that shows mostly similar effects of sex hormone treatment on tested transcripts and suggests that the concentration of sex hormones, rather than the sex origin of the myotubes, largely drives transcriptional activity in muscle cells. Based on the studied transcriptional response to increasing concentrations of estradiol, progesterone and testosterone in this study, we found minor effects of estradiol and progesterone while testosterone appeared to be the main driver of sex-specific differences that were also found in the baseline muscle biopsies of the same individuals. Only testosterone induced transcriptomic changes at a concentration resembling that found in serum of males and the presence of testosterone during myotube differentiation was able to mimic observed sex-based differences in vivo by affecting the transcription of *MYL3, MYH6, MYBPH* and *MYBPC2.* Other fiber type-associated sex dimorphisms like *MYH1/2/3/4, LDHA, LDHB, IRX3, GPD1, ENO3,* and *PLIN2* could not be evoked by sex hormonal treatment of myotubes. The transcriptional response of myotubes to testosterone, estradiol or progesterone can be different to the *in vivo* response of myofibers due to altered receptor densities and abundance of signal transducing proteins. The obvious sex dimorphism in the expression in histone demethylases located on X and Y chromosomes (*KDM5* and *KDM6* genes), has apparently no major impact on autosomal gene expression in myotubes.

Given the clear sex-specific differences of skeletal muscle on a multi-OMICs level, we anticipated a differential response to an individual defined and controlled endurance exercise bout. While other studies only reported the cumulative response over an extended period of training ^32,33,66^, we also analyzed skeletal muscle biopsies collected after the first acute exercise bout in our study. Indeed, the initial transcriptomic response in skeletal muscle showed clear differences between females and males. Females responded with upregulating transcripts essential for aerobic glycolysis, carbon-, pyruvate metabolism and TCA cycle, while the predominant response in males was associated with cellular and oxidative stress. We are not aware of similar studies comparing the acute transcriptional response of females and males to an exercise bout, but higher levels of circulating markers of muscle damage after acute bouts of endurance or strength training were regularly observed in males as well as higher levels of oxidative stress markers ^67–69^. These differences are often attributed to the protective effects of estradiol in females. Post-exercise leakage of muscle proteins into circulation such as myoglobin or creatine kinase is largely explained by exercise-induced oxidative damage and sarcolemma membrane destabilization. Estradiol treatment of ovariectomized rodents attenuated the release of creatine kinase and muscular immune cell infiltration ^70^ Estradiol localized to mitochondria can reduce mitochondrial emission of hydro peroxides and support maintenance of cellular redox state ^71^. Estradiol therapy has been associated to increased anti-oxidative capacities ^72,73^. Both, estradiol receptor-mediated and receptor-independent mechanisms are discussed to protect skeletal muscle membranes from exercise-induced damage ^74^, the latter one due to the structural properties of estradiol allowing its intercalation in membranes and to quench radicals ^71,73^. Our data corroborated this sexual dimorphism in the acute response to one exercise bout by the different regulation of transcripts known to be activated in response to cellular and oxidative stress. One caveat when comparing the acute response to an individualized exercise bout is that males in order to achieve the same individual exercise intensity had, in average, to produce more watts and greater muscle mass was engaged. While the differential regulation of the stress-responsive transcripts does not significantly correlate with the achieved performance in [W], the effect of muscle mass on the plasma concentrations of muscle-specific proteins measured after acute exercise cannot be neglected.

The comparison of the proteomic response after 8 weeks of training between females and males does not indicate any harmful consequences of the initial stress response of male muscle. Instead, our data show that adaptations to 8 weeks of endurance training were similar. Both females and males upregulated proteins and key enzymes of the TCA cycle, respiratory chain and β-oxidation and ATP synthesis. The results are well in line with our recently published data reporting that the increase in mitochondrial respiration assessed by high resolution respirometry in myofibers isolated from skeletal muscle biopsies of the subjects did not differ between sexes ^36^. We also did not observe differences in mitochondrial respiration before the intervention, quite in contrast to the respiration in subcutaneous adipose tissue analyzed in the same subjects, which was higher in females compared to males. Thus, the higher content of oxidative fibers in female muscle is not reflected by a higher mitochondrial respiration in albeit randomly selected myofibers. We analyzed the myofibers after respirometry for the abundance of MYH1,2 and 7 to estimate the fiber type proportion, but adjusting to differences in the MYH protein content did not give additional hints to a different mitochondrial oxidative capacity of female and male skeletal muscle when assessed in isolated myofibers.

Some key enzymes of acetyl-CoA metabolism showed a divergent regulation in females and males after the 8 weeks of training. The reduction of the cytosolic ACSS2 but not the mitochondrial ACSS1 enzyme in female muscle after training may favor reduced lipid synthesis and pave the way for enhanced supply of acetyl-CoA for citrate synthesis and fueling the TCA cycle. Interestingly, HMGCS2 was found to be increased only in female muscle after the 8 weeks training. HMGCS2 catalyzes the second and rate-limiting step of ketogenesis. Ketone production is not considered to play a major role in skeletal muscle but recently, β-hydroxybutyrate was described as protective factor for skeletal muscle by preventing muscle mass loss, mitochondrial impairments and functional decline ^75^. The intramuscular production of β-hydroxybutyrate can be a yet underestimated contributor to this mechanism, in addition to local ketone bodies serving as alternative fuel for skeletal muscle during exercise. The regulation of enzymes of acetyl-CoA synthesis and metabolism after 8 weeks of endurance training can be an adaptive process particularly in females, shifting acetyl-CoA from lipid synthesis to mitochondrial TCA cycle and ketogenesis thereby supporting increased fuel oxidation and protective mechanisms.

All described proteomic adaptations in female and male muscle are well in line with increased aerobic endurance exercise performance. This process is particularly visible in male muscle with a reduction in fast-twitch type 2 fiber-specific proteins and an overall reduction of sex-specific differences in enzymes of glycogen degradation and glycolysis suggesting a fiber type shift from glycolytic fast-twitch fibers to more oxidative fibers. An upregulation of proteins with a preferential expression in type 1 fibers was found in both sexes and was not more evident in male compared to female muscle. When individuals who perform endurance training on a regular basis were compared to sedentary ones ^35^, the proteomic profile of endurance-trained individuals showed a significant higher abundance of mitochondrial proteins, proteins involved in oxidative phosphorylation, the TCA cycle, and fatty acid metabolism. These protein signatures are consistent with the characteristics of type 1 (slow-twitch) muscle fibers, and suggests that long-term endurance training promotes adaptations typically found in type 1 fibers, enhancing aerobic capacity and energy efficiency in these muscles. Notably, when females and males in the endurance trained group were compared, only one protein was found to be different, in contrast to 30 proteins in the control groups ^35^. The marginal sex differences in the proteome of the endurance trained group are well in line with our observation of an equalization of baseline differences after 8 weeks of endurance training. Transcriptomic profiling of endurance trained female and male skeletal muscle and comparison with sedentary controls also revealed that sex differences are reduced in the endurance trained groups ^76^. Overall, the results of our study and of others hint towards the idea that regular endurance training might, over years, reduce or even eliminate sex-based proteomic differences in skeletal muscle, leading to a more similar proteomic profile across males and females. Our data indicate that this kind of harmonization already starts after 8 weeks of endurance training. Whether strength training would also result in an equalization of sex-specific differences in the skeletal muscle’s proteome presumably toward a more glycolytic fast-twitch fiber type is currently unclear.

Our study has limitations. As sex-based differences were not the main focus in the initial study design, the number of female and male participants is unequal. We were aware of this limitation during our data analysis, and we often specifically focused on what we observed as sex-specific differences in males as the underrepresented group in this study to avoid confounding effects by differences in number. The menstrual cycle was not tightly tracked in female participants, two used contraceptives and one subjects was post-menopausal. However, none of them was an outlier in PCA analyses of the OMICs data. Although hypermethylation in the skeletal muscle of female participants was observed across all chromosomes, the comparison of monoallelic (in men) and biallelic (in women) DNA methylation levels of CpG sites located on the X chromosome remains debatable ^77^. Furthermore, our data were obtained in a cohort with overweight or obesity and low cardiorespiratory fitness due to a sedentary lifestyle. It remains to be shown whether the results of the initial differences and in the adaptation to the 8 weeks of training can be translated to other cohorts differing in age, physical fitness and metabolic parameters.

In summary, we provide a first-of-it’s-kind multi-OMICs characterization of skeletal muscle. The pronounced sex-specific differences in transcripts and proteins with a preferential expression in type 1 or type 2 fibers mirror the functional differences in glucose and lipid metabolism and are deeply rooted in the epigenome.

Sex has also an impact on the response to exercise. Our data hint towards a potential protective effect of estrogen as we provide evidence on the transcriptional level that males experience higher oxidative and cellular stress in their muscles when cycling at the same individual intensity as females. The reduction in type 2 fiber-specific proteins specifically in males after only 8 weeks of training contributes to the endurance-trained proteomic profile in both sexes and underlines the high plasticity of skeletal muscle to adapt to repeated endurance exercise loads towards a metabolically beneficial phenotype independent of sex.

## Data availability

The transcriptomic data used in this study is available via the GEO database at NCBI (GSE224310). The epigenomic data used in this study is available via the GEO database at NCBI (GSE244359). The proteomic data of this study is available via the PRIDE database with identifier PXD058700. Further data that support the findings of this study are available from the corresponding author upon request.

## Acknowledgements

The authors thank all biopsy donors. The authors are grateful for the excellent technical support provided by Kolja Leffek from the University Hospital Tübingen, Tübingen, Germany, and acknowledge the technical support of Core Facility Metabolomics and Proteomics at Helmholtz Munich.

---

**Figure S 1.**
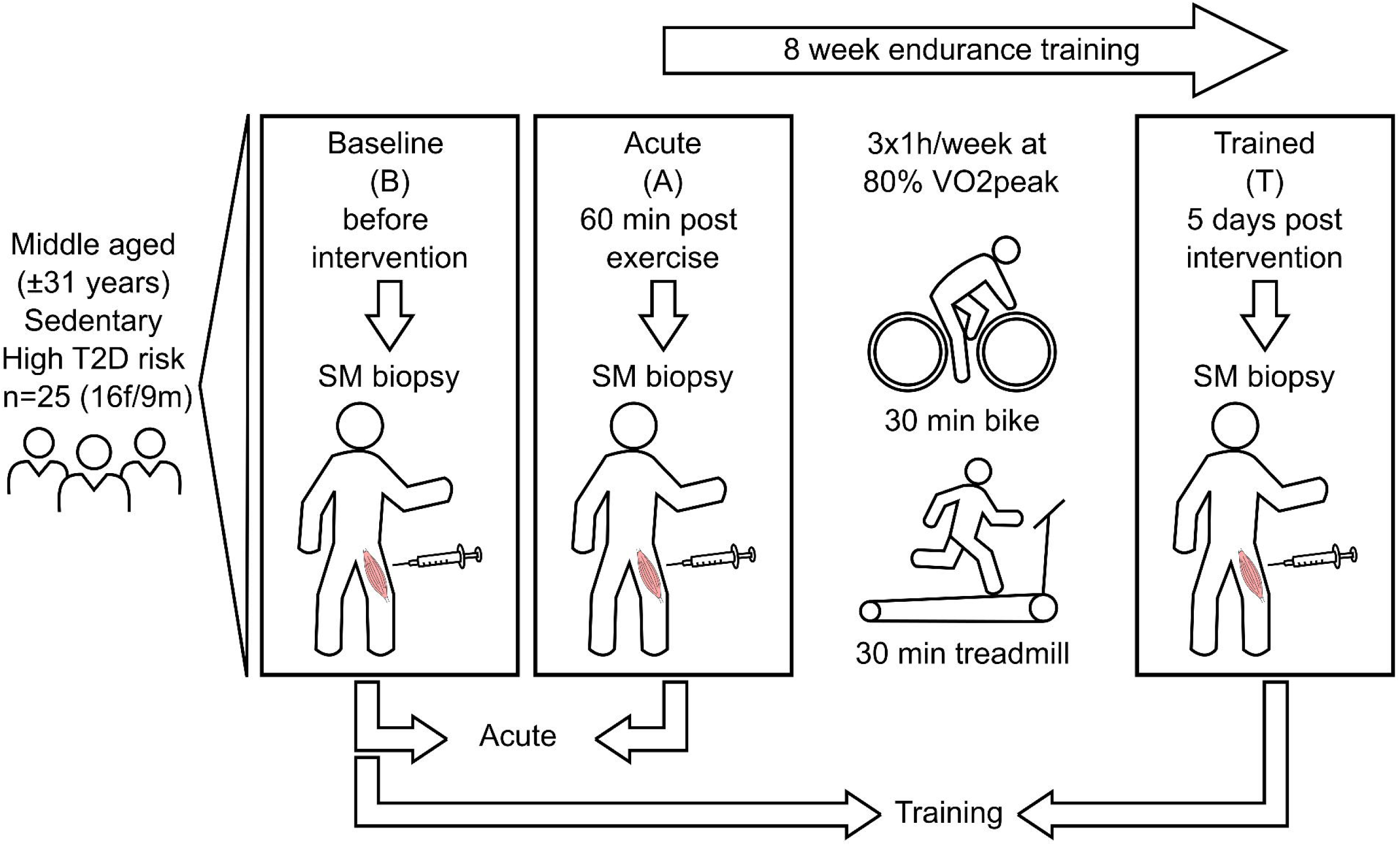
Timeline of the exercise intervention study Middle aged participants with a sedentary lifestyle and overweight or obesity (n=25, 16f/9m) underwent an 8-week supervised endurance exercise intervention consisting of 30min on ergometer and 30min on a treadmill at 80% VO2peak. Muscle biopsies were obtained at baseline (B) before exercise, 60min after the first acute ergometer exercise in an untrained state (A) and 5 days after the last exercise session after 8 weeks of training (T) in a rested state.

**Figure S 2.**
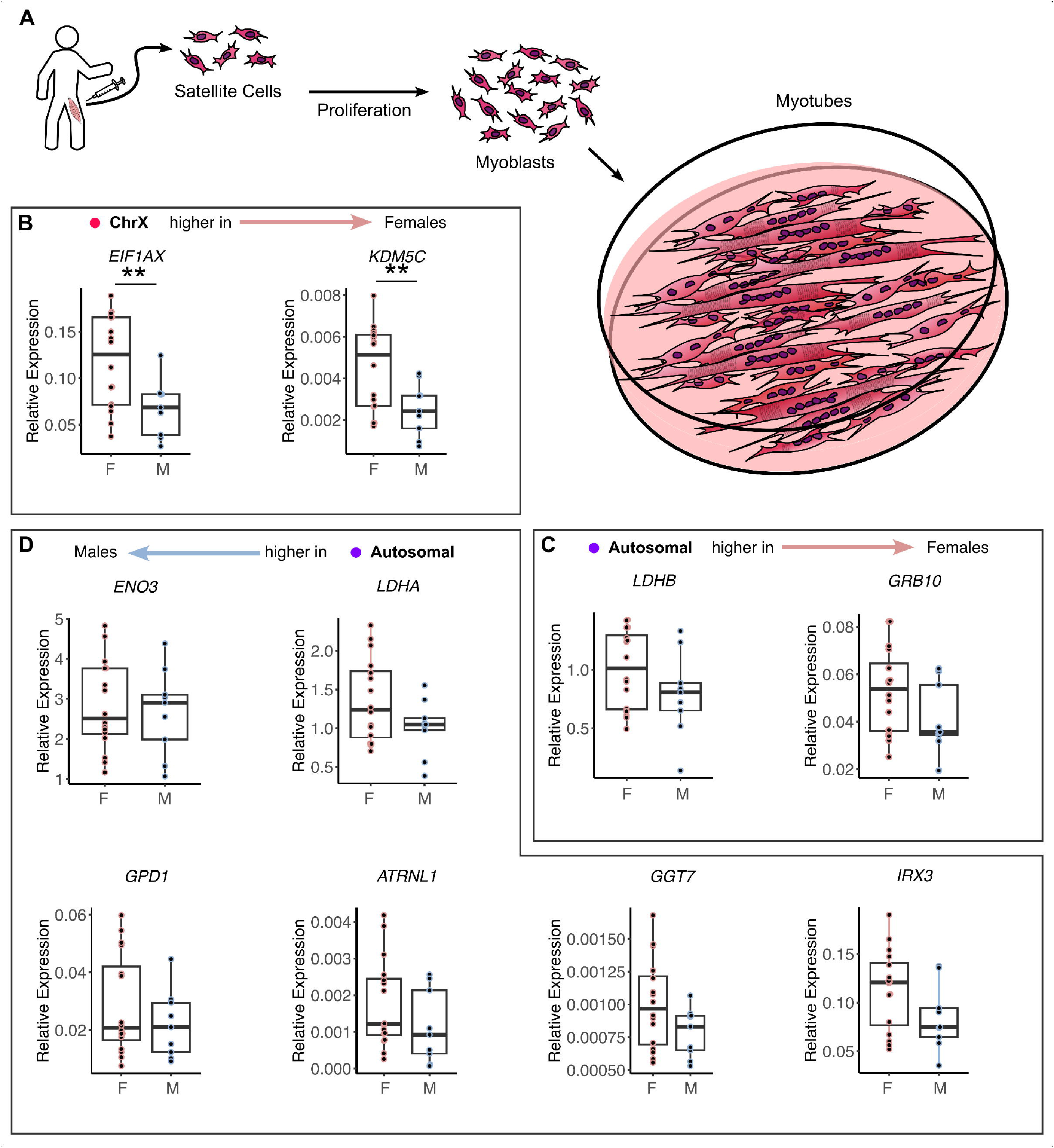
Conserved sex-specific differences in myotubes in vitro A) Satellite cells were isolated from skeletal muscle biopsies, myoblasts were cultured and differentiated into myotubes for 7 days. Expression of B) X-, Y- chromosomal and C/D) autosomal genes is shown for female (F) and male (M) myotubes. Arrows indicate sex-specific differential expression in vivo in skeletal muscle biopsies. Significant differences were assessed using one-way ANOVA LSD, ** p<0.01, *** p<0.001, n=25 (16f/9m).

**Figure S 3.**
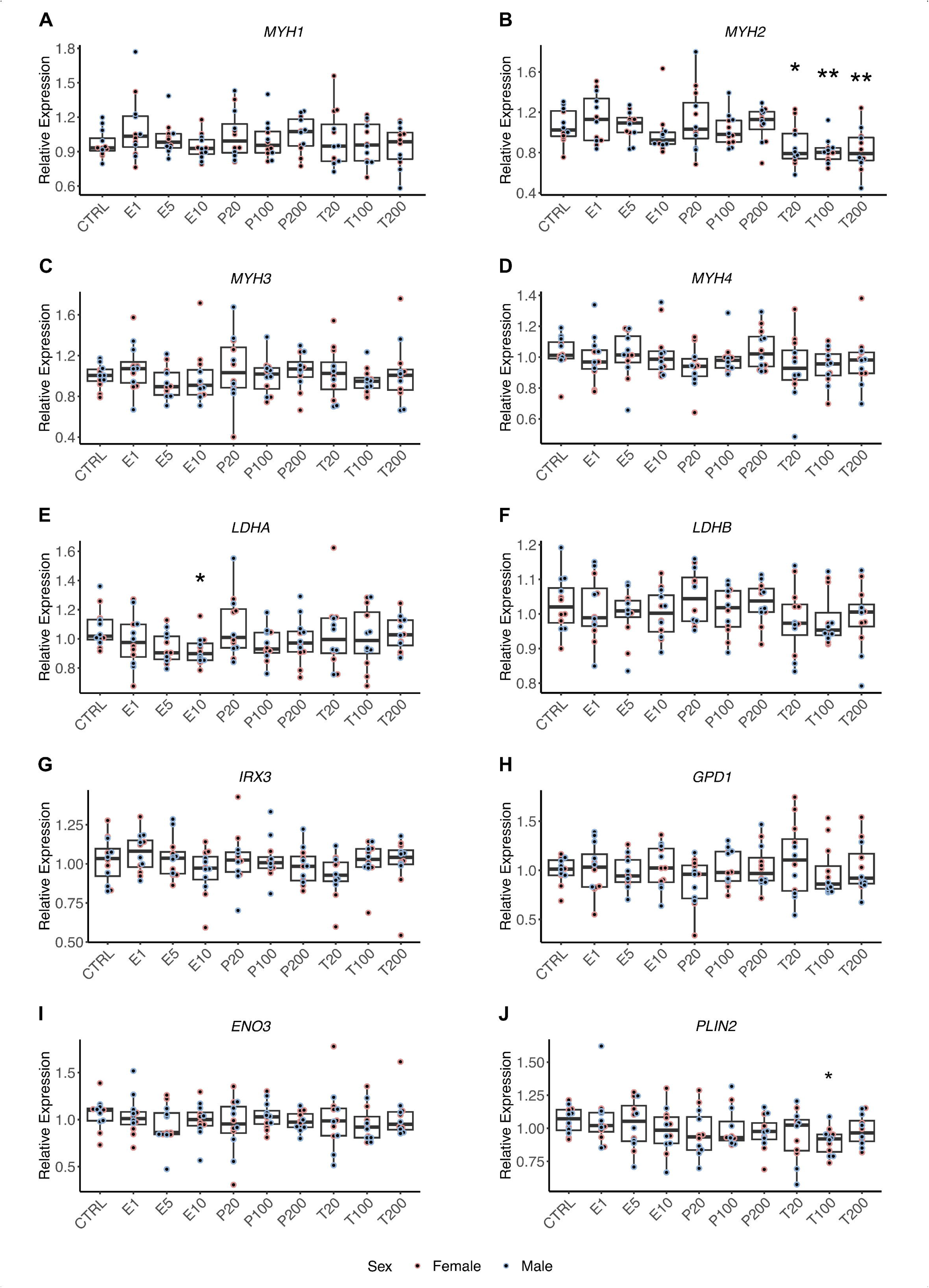
Sex hormone-specific transcriptional regulation in myotubes in vitro A) Myoblasts derived from 6 female and 6 male donors were cultured and differentiated into myotubes for 7 days. During differentiation, myotubes were treated with estradiol (E) (1/5/10nM), progesterone (P) (20/100/200nM) or testosterone (T) (20/100/200nM) or left untreated (CTRL). Expression of mRNA was measured for A) MYH1, B) MYH2, C) MYH3, D) MYH4, E) LDHA, F) LDHB, G) IRX3, H) GPD1, I) ENO3, J) PLIN2 in myotubes after hormonal treatment, respectively. Statistical significance was determined by using one-way ANOVA Bonferroni, n=12 (6f/6m). Red dots, female donor; blue dots; male donor.

**Figure S 4.**
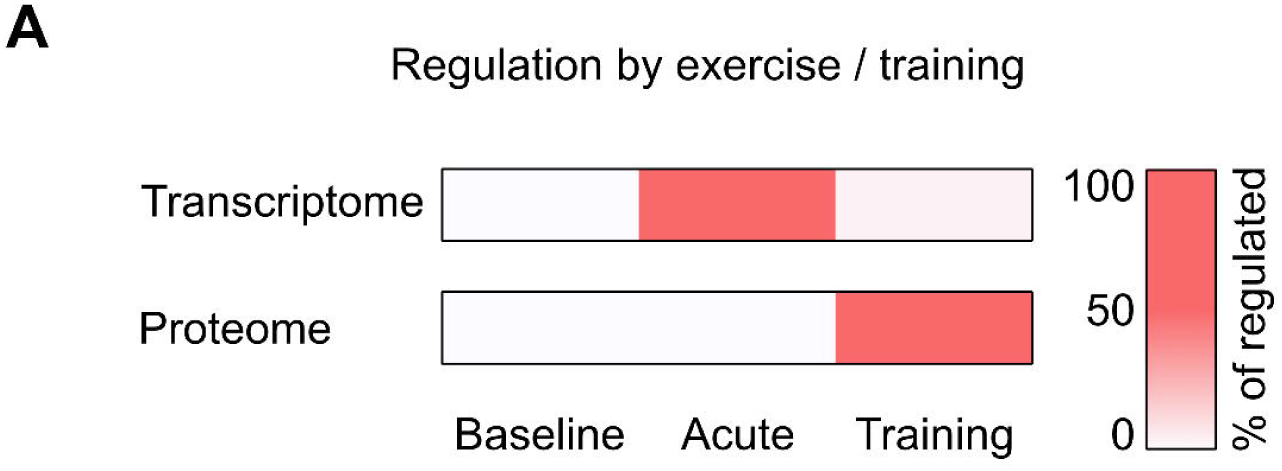
Transcriptomic and proteomic responses to acute exercise and training A) Heatmap depicting the percentage of all transcripts or proteins regulated by acute exercise and training. Out of all transcripts regulated by exercise and training, 98% were regulated in response to acute exercise and 4% after training. Out of all proteins regulated by exercise and training, 1% were regulated in response to acute exercise and 100% after training. Statistical significance was determined by limma t-test BH, p < 0.05, n=25 (16f/9m).

**Supplementary Table 1.** Correlation of Transcription and Performance.

| Pearson Correlation with Performance [W] |  |  |
| --- | --- | --- |
|  | r | p-Value |
| *ATF3* | -0.040 | 0.848 |
| *JUN* | 0.053 | 0.800 |
| *STK39* | 0.246 | 0.235 |
| *SLC39A14* | 0.235 | 0.258 |
| *HMOX* | 0.234 | 0.259 |
| *MT1A* | 0.119 | 0.570 |
| *MT1B* | 0.022 | 0.916 |

Supplementary Data Table 1 Excel file containing epigenomic, transcriptomic and proteomic data analysis of the study

